# A metabolic CRISPR-Cas9 screen in Chinese hamster ovary cells identifies glutamine-sensitive genes

**DOI:** 10.1101/2020.05.07.081604

**Authors:** Karen Julie la Cour Karottki, Hooman Hefzi, Songyuan Li, Lasse Ebdrup Pedersen, Philipp Spahn, Chintan Joshi, David Ruckerbauer, Juan Hernandez Bort, Alex Thomas, Jae Seong Lee, Nicole Borth, Gyun Min Lee, Helene Faustrup Kildegaard, Nathan E. Lewis

**Affiliations:** The Novo Nordisk Foundation Center For Biosustainability, Technical University Of Denmark, Denmark; The Novo Nordisk Foundation Center For Biosustainability At The University Of California, San Diego, USA; Department Of Biological Sciences, Kaist, 291 Daehak-Ro, Yuseong-Gu, Daejeon 305-701, Republic Of Korea; Department of Pediatrics, University of California, San Diego, USA; Department of Bioengineering, University of California, San Diego, USA; Austrian Centre of Industrial Biotechnology, Vienna, Austria; University of Natural Resources and Life Sciences, Vienna, Austria; Department of Molecular Science and Technology, Ajou University, Suwon 16499, Republic of Korea

**Keywords:** CHO, CRISPR pooled screen, glutamine, metabolism^1^

## Abstract

Over the past decades, optimization of media formulation and feeding strategies have fueled a many-fold improvement in CHO-based biopharmaceutical production. While Design of Experiments (DOE) and media screens have led to many advances, genome editing offers another avenue for enhancing cell metabolism and bioproduction. However the complexity of metabolism, involving thousands of genes, makes it unclear which engineering strategies will result in desired traits. Here we developed a comprehensive pooled CRISPR screen for CHO cell metabolism, including ∼16,000 gRNAs against ∼2500 metabolic enzymes and regulators. We demonstrated the value of this screen by identifying a glutamine response network in CHO cells. Glutamine is particularly important since it is often substantially over-fed to drive increased TCA cycle flux but can lead to accumulation of toxic ammonia. Within the glutamine-response network, the deletion of a novel and poorly characterized lipase, *Abhd11*, was found to substantially increase growth in glutamine-free media by altering the regulation of the TCA cycle. Thus, the screen provides an invaluable targeted platform to comprehensively study genes involved in any metabolic trait.

---

Chinese hamster ovary (CHO) cells are the most commonly used mammalian cells for biotherapeutic protein production and serve as the expression system of choice for the leading biologics^1^. Consequently, improving product quality and decreasing manufacturing costs in CHO is of great interest to the biopharmaceutical industry. Since their first use in the late 1980s, final product titers from CHO cells have improved more than 50-fold, largely through media and bioprocess optimization^2^. Although effective, these empirical approaches are highly variable, demand extensive labor, time, and resources, and may not translate directly to new clones.

All biological processes that lead to protein production depend on metabolic building blocks. Although CHO cell media are complex owing to their nutritional demands^3^ the two main nutrients consumed are glucose and glutamine. These are often taken up in excess of the cells growth needs^4^ leading to increased by-product formation of lactate and ammonia, respectively, which are the two primary byproducts negatively affecting cell growth, production and product quality^5–8^. The complexity and incomplete understanding of metabolism, along with unique idiosyncrasies of individual CHO clones have stymied the optimization of their metabolism. However, the release of CHO and Chinese hamster genome sequences^9–11^ and improved systems biology approaches^3,12^ have laid the groundwork for a new era of targeted CHO cell line development, but the question of the best way to discover and engineer targets remains open.

Several techniques can be used to knock out genes in CHO cells, such as zinc finger nucleases (ZFNs)^13^, transcription activator-like effector nucleases (TALENs)^14^, and Clustered Regularly Interspaced Short Palindromic Repeats (CRISPR). However, since the best genes to knock out are often unclear, given >20,000 genes in the CHO genome, efficient, high-throughput methods are needed to identify optimal genetic modifications. Although RNA interference (RNAi) screening has been useful for identifying gene knockdowns^15^ providing a desired trait in CHO cells^16^, the inability to achieve full knockout, a significant amount of off-target effects^17^, and inconsistent results has limited their use^18^. On the other hand, CRISPR-Cas9 can also be used for large-scale pooled screening while avoiding some of the pitfalls in RNAi screens^19^. The method has been established in several cell lines and organisms, mainly mouse and human, increasing the robustness for the next generation of forward genetic screening methods^19–23^.

With the intent of generating a platform for gaining insight into CHO cell metabolism, we present a large-scale CHO-specific CRISPR-Cas9 knockout screen in CHO cells. We generated a gRNA library targeting genes for enzymes and regulators involved in CHO cell metabolism. We deployed CRISPR-Cas9 knockout screening against an industrially relevant selection pressure, glutamine deprivation, and identified a network of genes regulating growth in response to glutamine concentration. We highlight one gene for a novel and poorly characterized lipase, *Abhd11*, which, upon deletion, we found to substantially increase growth glutamine-free media by altering the regulation of the TCA cycle.

## Results

### Establishing a CRISPR knockout library in CHO cells

We first generated CHO-S cell lines constitutively expressing Cas9 (CHO-S^Cas9^) via G418 selection followed by single cell sorting and expansion to obtain clonal populations for subsequent gRNA library transduction. We validated the functionality of Cas9 in the clonal cell lines by transfecting CHO-S^Cas9^ with a gRNA targeting *Mgat1* and quantifying the cleavage efficiency by indel analysis of the target region (Supplementary Table S1). To generate the CRISPR knockout library, we designed a large CHO-specific gRNA library containing 1-10 gRNAs/gene for genes encoding enzymes and regulators of CHO metabolism. Genes selected for inclusion were obtained from the genome scale metabolic model of CHO^3^, metabolism-associated GO terms, and transcription factors that regulate the aforementioned genes (based on annotation from Ingenuity Pathway Analysis^24^). The library consists of 15,654 gRNAs against 2,599 genes (1,765 genes from the model, 782 from GO terms, and 52 transcription factors)(Supplementary Datafile 1). gRNAs were synthesized by CustomArray Inc. and subsequently packaged into lentiviruses. CHO-S^Cas9^ cells were then transduced with the gRNA library at low multiplicity of infection (MOI) (Supplementary Methods and Results) to ensure only a single gRNA integration event per cell, generating a CHO CRISPR knockout library for use in pooled screening (overview in Figure 1).

**Figure 1.**
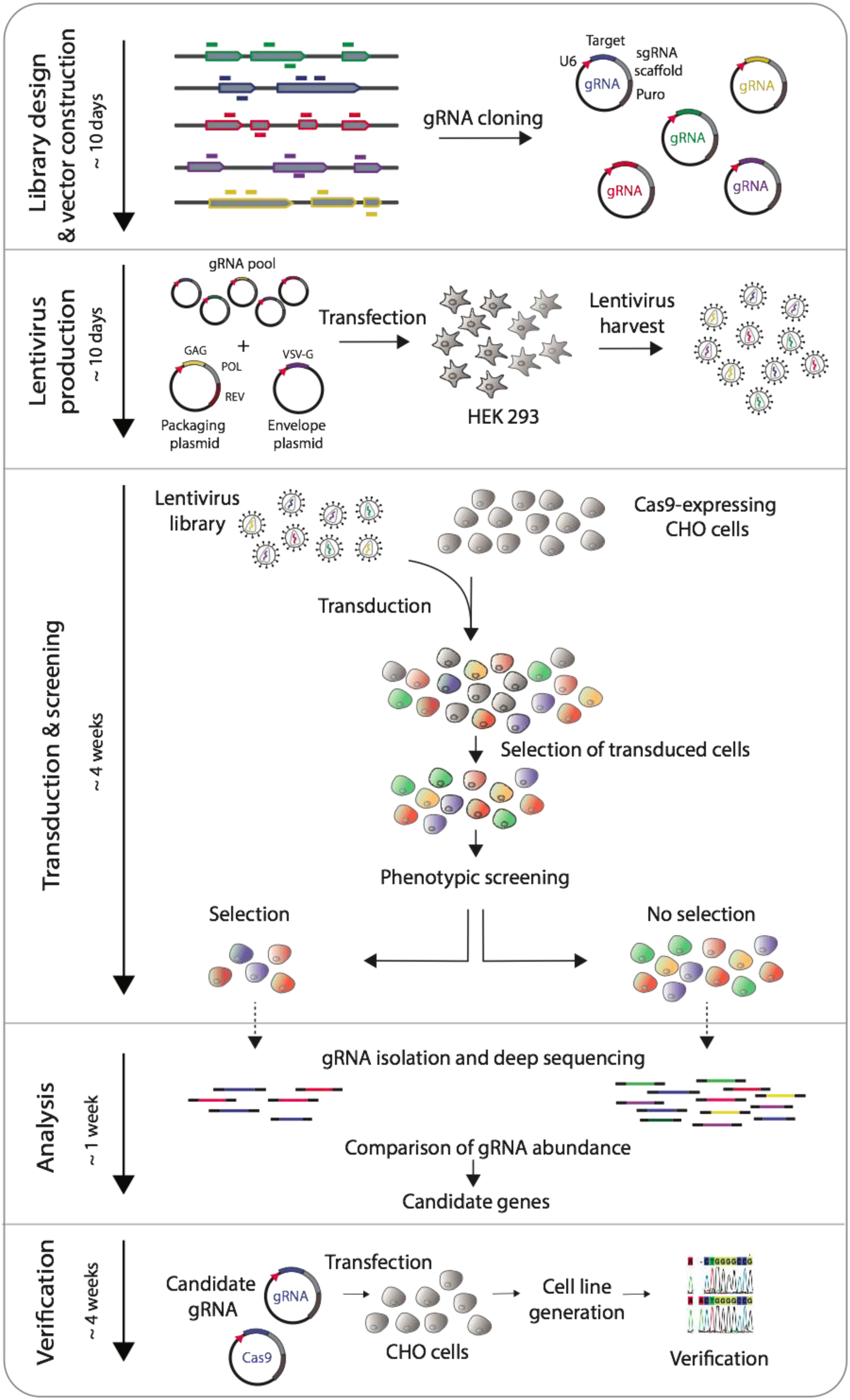
Screening overview. gRNAs are computationally designed to target the genes of interest, then synthesized and cloned into gRNA scaffold containing vectors. HEK cells are transfected with packaging vectors and gRNA vectors to generate a pool of viruses containing all the gRNA vectors. After harvest, the pooled library is used to transduce Cas9-expressing CHO cells at a low MOI to ensure a single integration event per cell. Cells positive for gRNA integration are selected for with antibiotics before undergoing a phenotypic screen. Genomic DNA is extracted from the collected cells and gRNA presence is compared between samples. Enriched or depleted gRNAs are ranked and candidate genes are phenotypically validated.

### Glutamine screening

Glutamine is key to cell function and thus an important media component for animal cell culture media formulations^25^. However, glutamine is often oversupplied, and its catabolism produces ammonia, a toxic byproduct that negatively impacts cell growth, production, and product quality^5,26–28^. Understandably, it is of interest to identify engineering strategies that permit improved cell behavior in glutamine-free conditions. We thus screened the CHO CRISPR knockout library cells for growth in media with and without glutamine for fourteen days. The cells were passaged every three days (growth profile in Supplementary Figure S1) and 30 × 10^6^ cells were collected at the beginning and the end of the screen for analysis to ensure adequate coverage.

### The gRNA library is well represented at the start of screening

To ensure that all possible gene knockouts are screened it is important to verify that the gRNA library is well represented at the beginning of the screen. We therefore sampled the cells just prior to glutamine deprivation (T0) and sequenced the gRNAs present in the starting cell pool. From the entire library, only 2 genes (<0.1%) and 638 gRNAs (<4%) were absent at the initial time point. In all samples, median-normalized gRNA and gene sequencing depth was greater than 35 and 360 CPM (counts per million), respectively (Figure 2). Thus, the majority of the library was well represented before the CRISPR knockout library was subjected to screening.

**Figure 2.**
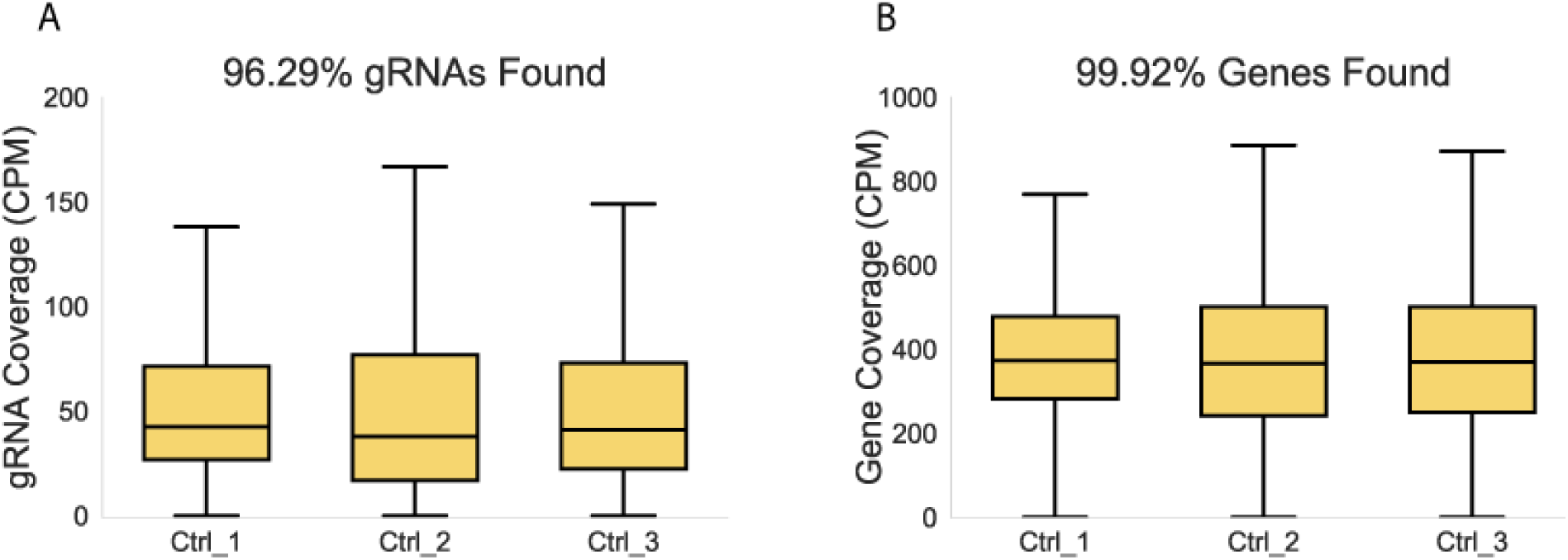
Screen verification. A) Read count per gRNA. B) Total read count per gene (summed over all gRNAs). Shown are normalized read counts (counts per million/CPM) for three replicate experiments prior to starting selection. Outliers not displayed.

### Glutamine screening reveals expected and novel targets

To identify gRNAs impacting CHO cell growth in glutamine free media, we analyzed gRNA enrichment and depletion between samples grown for fourteen days in media with and without glutamine. As expected, the absence of glutamine does not display a strong selection pressure (Supplementary Figure S2), consistent with the ability of CHO cells to grow slowly in the absence of glutamine due to low levels of endogenous glutamine synthetase expression^29^. We found 20 genes (Figure 3) that were significantly enriched or depleted in all replicates. As expected, *Glul* (glutamine synthetase) gRNAs showed significant depletion in cells grown without glutamine, consistent with its role as the enzyme responsible for *de novo* glutamine synthesis. Similarly, significant enrichment of *Gls* (glutaminase) gRNAs was observed, consistent with protection of the intracellular glutamine pool from undesirable catabolism when glutamine is not readily available.

**Figure 3.**
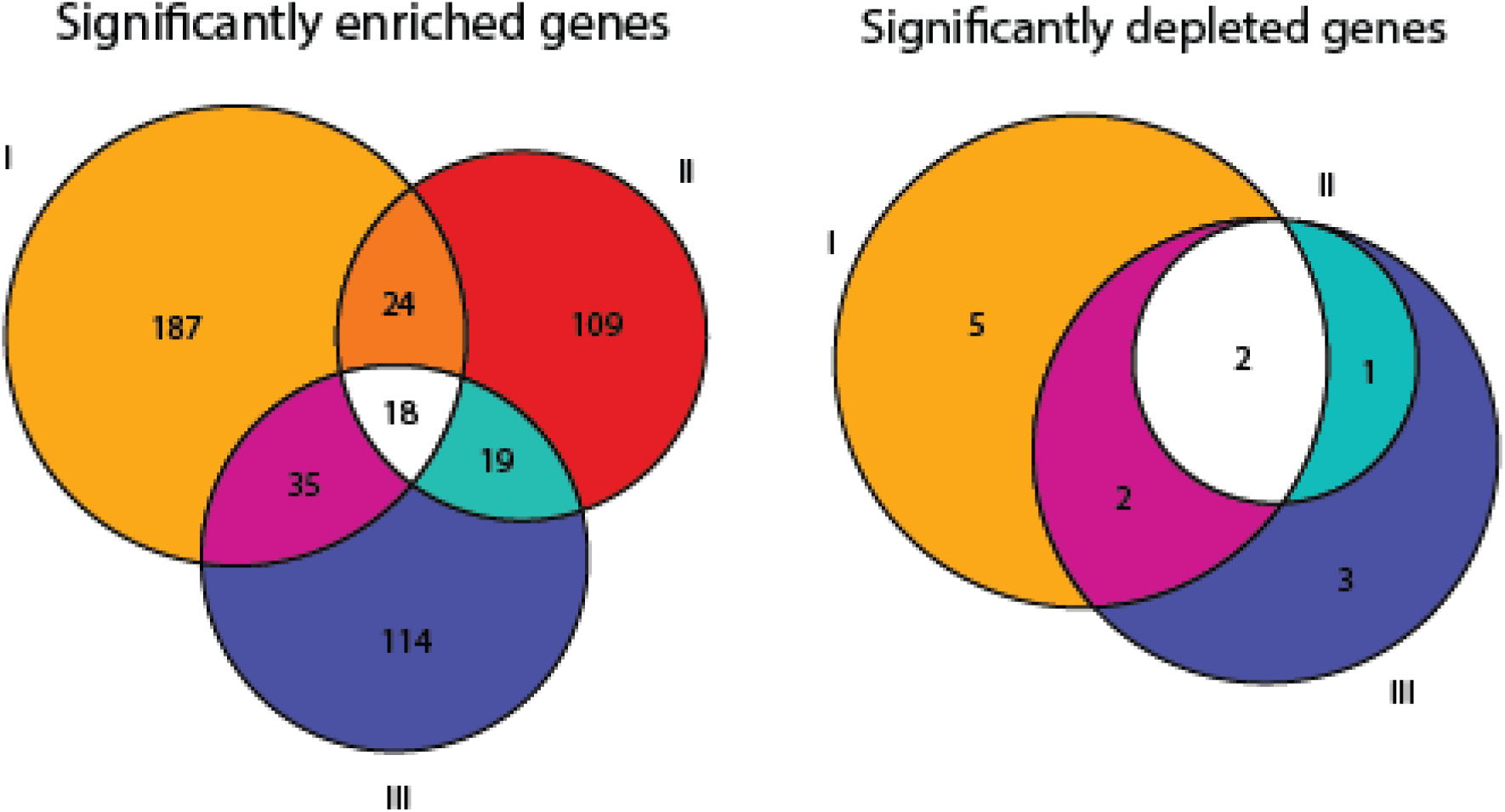
Significantly enriched and depleted genes following glutamine selection. Three glutamine screens of the knockout library were carried out and the significantly depleted and enriched genes from each replicate are shown. While there was variability between replicates (I-III), eighteen significantly enriched genes and two significantly depleted genes were commonly observed in all experiments.

### Disruption of *Abhd11* is conditionally beneficial dependent on presence of glutamine

We found the strongest and most consistent gRNA enrichment in cells grown without glutamine was a poorly characterized putative lipase, *Abhd11*. We subsequently generated clonal *Abhd11* knockout cell lines using CRISPR-Cas9 and assessed their growth in media with and without glutamine. In accordance with the screen, knocking out *Abhd11* substantially improved growth in glutamine-free medium (Figure 4A) but also depressed growth in glutamine containing medium compared to control cells (Figure 4B).

**Figure 4.**
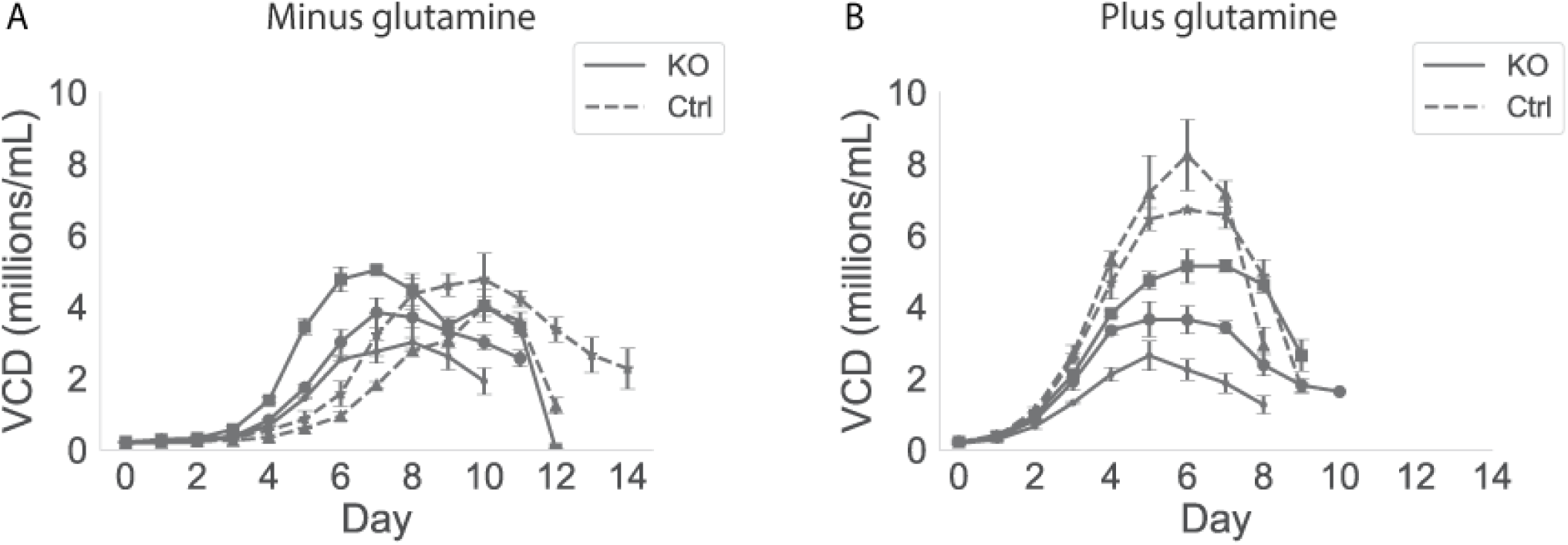
Growth curves for *Abhd11* knockout and control cell lines in batch culture in media without and with glutamine. Growth curves for three *Abhd11* knockout (KO) and two control (Ctrl) cell lines grown in three replicates in media without glutamine (A) and with glutamine (B). Viable cell density (VCD) was measured every day over a period of 14 days.

*Abhd11* has been poorly studied and is currently annotated as a putative lipase. However, recent work reports that Abhd11 associates with the alpha-ketoglutarate dehydrogenase complex (αkgdhc) and prevents its de-lipoylation^30^ (a crucial cofactor for its activity). The *Abhd11* knockout would thus be expected to decrease αkgdhc activity. The benefit of the knockout in glutamine-free (and detriment in glutamine replete) conditions is congruous with this mechanism. In the presence of glutamine, wildtype cells fuel the TCA cycle heavily via glutaminolysis^31^, without Abhd11, αkgdhc activity would be attenuated and entry of glutamine to the TCA cycle would be stunted. Consistent with this, we observe drastically increased glutamate secretion in KO cells when grown in media containing glutamine (Figure 5) and decreased glutamine uptake (KO cultures maintain >3 mM glutamine at all timepoints while wildtype cells consume all glutamine by day 5 or 6, data not shown).

**Figure 5.**
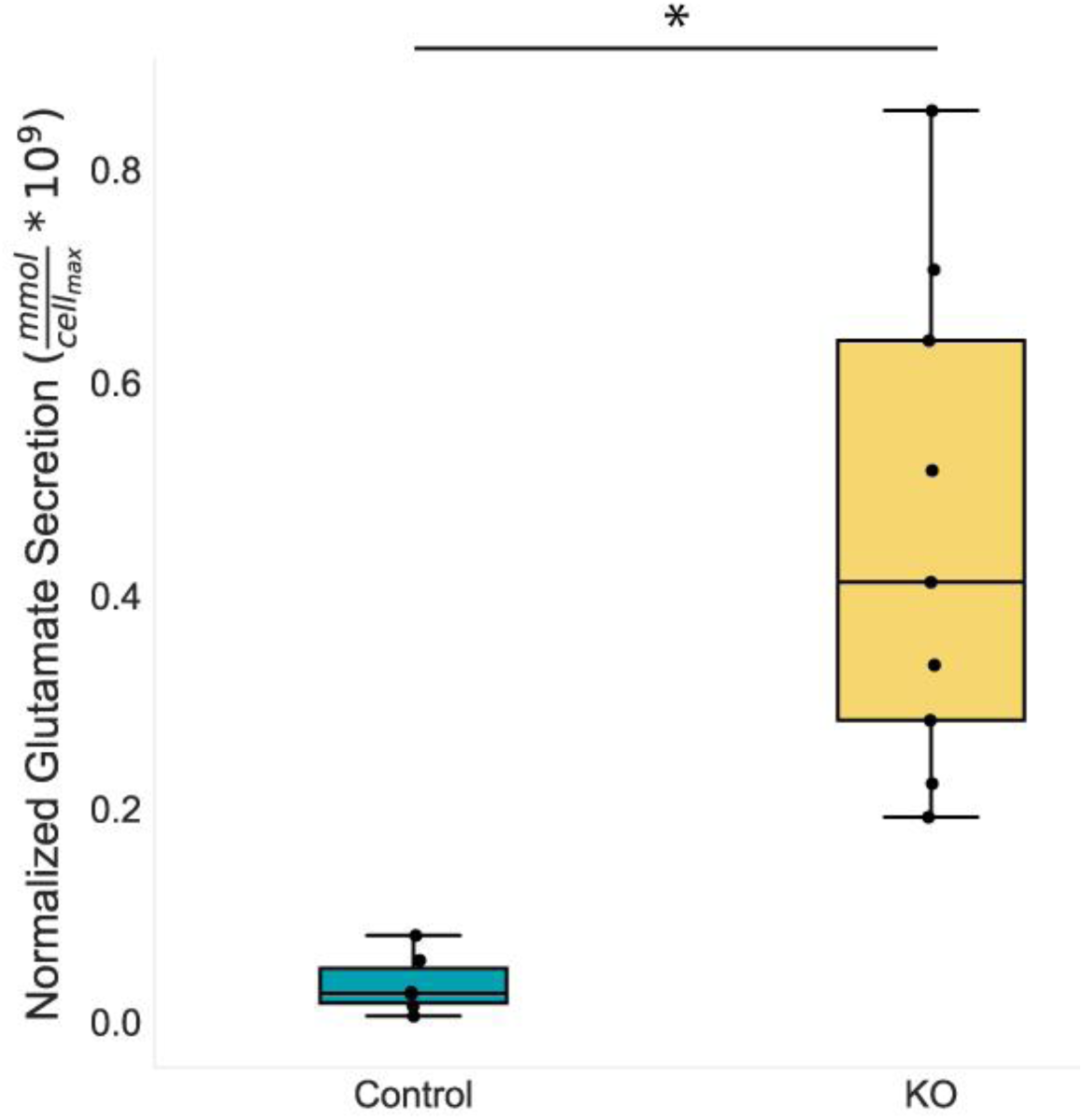
Impact of *Abhd11* knockout on glutamate secretion. Wild type (control) and knockout cells were grown in glutamine replete conditions. Glutamate secreted during the growth phase (e.g., until maximum VCD was reached) was normalized by the maximum VCD to approximate cell specific glutamate secretion. Knockout cells secreted significantly more glutamate than wildtype cells. * indicates a statistically significant difference (p<0.05) as calculated by a two-tailed Welch’s t-test.

In the absence of glutamine, the decrease in αkgdhc activity in knockout cells would act as an artificial bottleneck at alpha-ketoglutarate (αkg), forcing carbon away from the TCA cycle and into glutamine biosynthesis. Thus, control cells, with functional Abhd11, would consume αkg via αkgdhc to a greater extent than knockout cells, pulling away from *de novo* glutamine synthesis, which is essential for growth in glutamine-free medium. Indeed, when cells are adapted via stepwise decreases in glutamine levels and directed evolution^32^, cells adapt by decreasing their expression of *Abhd11* (Supplementary Datafile 2). An overview of the putative impact of Abhd11 on glutamine metabolism is shown in Figure 6.

**Figure 6.**
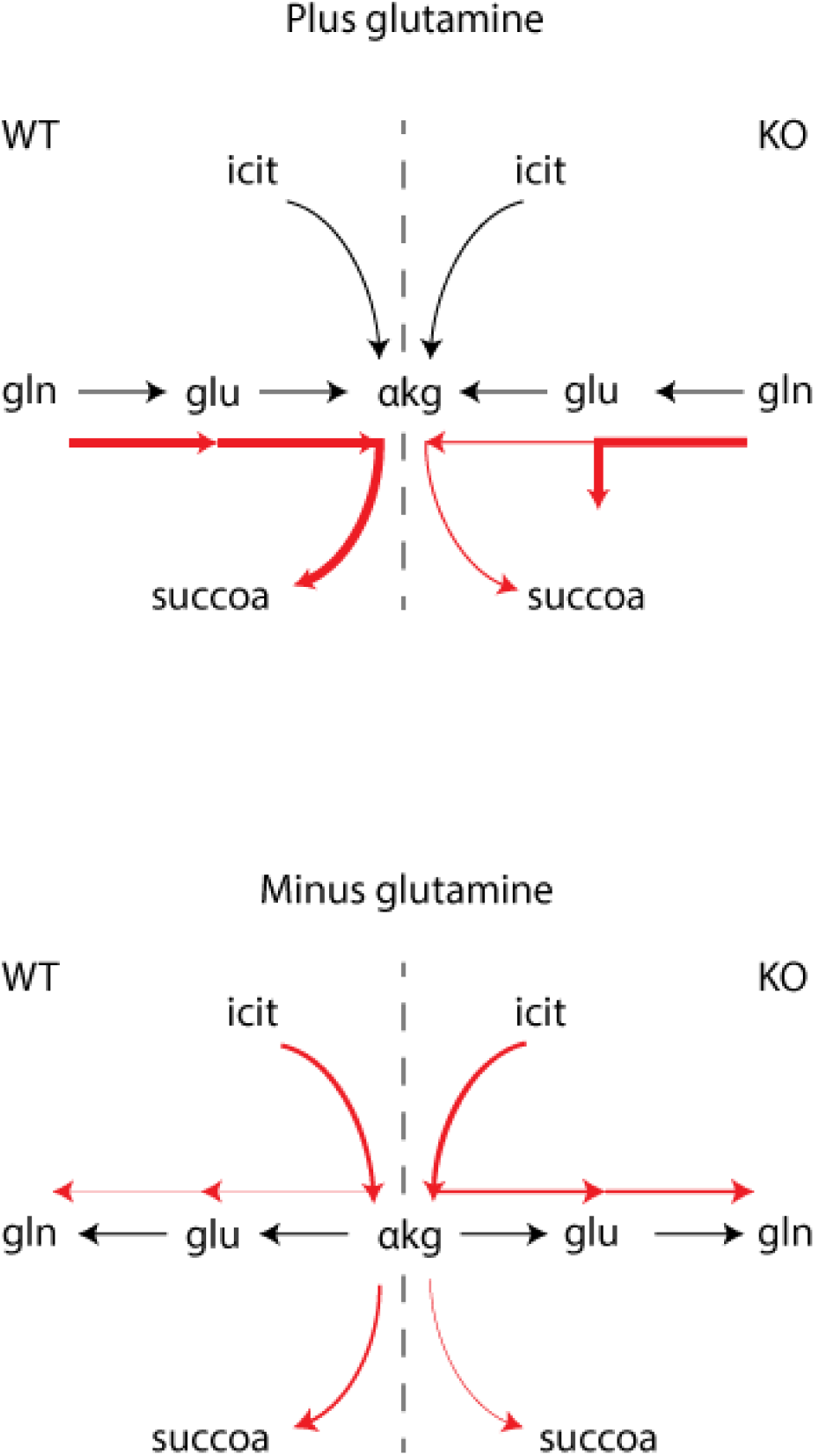
Putative mechanism of action for wild type (WT) and *Abhd11* knockout (KO) cells grown in media with or without glutamine. Abhd11 associates with and protects the αkgdhc. (a) In the presence of gln, cells fuel the TCA cycle through gln catabolism. In *Abhd11* KO cell lines, αkgdhc flux (and TCA cycle activity) is decreased, αkg and glu accumulate, and glu is secreted, leading to decreased growth for KO cells. (b) Without gln, the TCA cycle is largely fueled through glycolysis. In *Abhd11* KO cell lines, the decrease in αkgdhc activity leads to increased αkg, which permits increased flux to glu and *de novo* glutamine synthesis. With normal Abhd11 function, cells do not have this bottleneck and αkgdhc activity competes more strongly with gln biosynthesis, leading to decreased growth for WT cells. αkgdhc: alpha ketoglutarate complex, icit: isocitrate, αkg: alpha-ketoglutarate, succoa: succinyl coenzyme A, gln: glutamine, glu: glutamate.

To explore the relationship between *Abhd11* and glutamine metabolism, we further analyzed knockout and control cell lines and compared their transcriptomic profile when grown in media with and without glutamine (Supplementary Methods and Results).

## Discussion

As CHO cells are the primary workhorse for the production of biopharmaceuticals, significant time and effort has been invested towards producing optimal cell lines for growth, high protein titer, and good protein quality. Here, we present a high-throughput approach to identify novel targets for CHO cell line engineering. The objective was two-fold: first to establish a CHO-specific metabolic CRISPR-Cas9 knockout screening platform in CHO cells and second to use this platform to explore CHO cell metabolism using an industrially relevant screening setup. Glutamine is one of the major nutrients taken up by mammalian cells and plays an important role as an energy source in *in vitro* culture^25,33^. The fast consumption of glutamine results in accumulated ammonia in the medium, inhibiting cell growth, reducing productivity, and altering glycosylation patterns on heterologously expressed proteins^5,27,34^. While growth on glutamine-free media is possible, a significant decrease in growth rate is almost always observed^35^. It is therefore of interest to investigate genetic alterations that elicit a positive growth response to media lacking glutamine. We found several genes whose knockout resulted in a growth benefit in media without glutamine. Unsurprisingly, one of these genes was *Gls*, which codes for the primary glutamine-catabolizing enzyme. Many of the remaining targets found were novel with respect to their protective role in glutamine depletion in CHO cells and their roles in a biological context are a topic for further investigation. We chose to follow up on *Abhd11*, a gene with no clear link to glutamine metabolism that showed the most marked enrichment of gRNAs in cells grown under glutamine depleted compared to glutamine replete conditions. Our results are consistent with recent evidence linking Abhd11 with a protective role of αkgdhc in the TCA cycle^36^. We observed depressed growth of *Abhd11* knockout cells in glutamine containing media alongside glutamate accumulation in the media and lack of complete glutamine consumption. As glutamate (via glutaminolysis) is a major source of TCA cycle intermediates^31^, the secretion of glutamate (and assumed decrease in TCA cycle activity) is consistent with the observed reduced growth rate. Conversely, in glutamine free media, *Abhd11* knockout cells exhibited improved growth compared to the wild type cells. We postulate that the inhibition of α-ketoglutarate catabolism leads to accumulation of α-ketoglutarate and increases its availability for conversion to glutamate and subsequently to glutamine, leading to better growth.

High-throughput CRISPR-Cas9 screening presents a novel approach to conduct forward genetic engineering and can provide an abundance of knowledge in the study of genotype to phenotype relationships. Over recent years CRISPR-Cas9 screens have been applied to a variety of mammalian cell types to study biological function^20,21,37^. Since the publication of initial CRISPR-Cas9 screens, comprehensive reviews and extensive method articles have been published^38–40^. We show here that CRISPR screening techniques can be applied to the industrially relevant CHO cell line. This approach enables a wide array of studies in CHO cells by applying different screening conditions or exploiting the existing variations of the Cas protein, such as catalytically inactive Cas9 coupled to transcriptional activators and repressors, for activation or repression screens as has shown potential in other mammalian cells^39,41–45^. With continuous advances in CRISPR screen design and comprehensive annotation of the CHO cell genome these types of screens will enable a new era of targeted engineering to improve CHO cell phenotypes.

## Methods

### Plasmid design and construction

The GFP_2A_Cas9 plasmid was constructed as previously described^46^. A Cas9 expression vector for generation of a Cas9 expressing CHO cell line (from here on be referred to as CHO-S^Cas9^), was constructed by cloning the 2A peptide-linked Cas9 ORF from the GFP_2A_Cas9 expression vector^46^ into a pcDNA™3.1(+) vector (Thermo Fisher Scientific) between the HindIII and BamHI sites. The construct will from here on be referred to as pcCas9. gRNA vectors were constructed using Uracil-Specific Excision Reagent (USER) friendly cloning as previously described^47^. Plasmids were purified using NucleoBond Xtra Midi EF (Macherey-Nagel) according to manufacturer’s protocol. Target sequences and gRNA oligos are listed in Supplementary Table S2.

### Cell culture

CHO-S wild type cells from Life Technologies were cultivated in CD-CHO medium (Thermo Fisher Scientific) supplemented with 8 mM L-Glutamine and 2 µL/mL AntiClumping Agent (AC) (Thermo Fisher Scientific) in a humidified incubator at 37 °C, 5 % CO2 at 120 RPM shake in sterile Corning® Erlenmeyer culture flasks (Sigma-Aldrich) unless otherwise stated. Viable cell density (VCD) was measured using the NucleoCounter® NC200™ (Chemometec) utilizing fluorescent dyes acridine orange and 4’,6-diamidino-2phenylindole (DAPI) for the detection of total and dead cells. Cells were seeded at 0.3 × 10^6^ cells/mL every three days or 0.5 × 10^6^ cells every two days.

### Transfection and cell line generation

For all transfections, CHO-S wild type cells at a concentration of 1 × 10^6^ cells/mL in a six well plate (BD Biosciences) in AC free media were transfected with a total of 3.75 μg DNA using FreeStyleTM MAX reagent together with OptiPRO SFM medium (Life Technologies) according to the manufacturer’s instructions. For generation of CHO-S^Cas9^, CHO-S wild type cells were transfected with pcCas9. Stable cell pools were generated by seeding transfected cells at 0.2 × 10^6^ cells/mL in 3 mL selection media containing 500 µg/mL G418 (SigmaAldrich) in CELLSTAR® 6 well Advanced TC plates (Greiner Bio-one) two days post transfection. Medium was changed every four days during selection. After two weeks of selection, cells were detached and adapted to grow in suspension. The clonal cell lines were analysed by Celigo Cell Imaging Cytometer (Nexcelom Bioscience) based on the green fluorescence level using the mask (blue fluorescence representing individual cells stained with NucBlue™ Live ReadyProbes™ Reagent; Thermo Fisher Scientific) + target 1 (green fluorescence) application. For generating knockout cell lines of screen targets, CHO-S wild type cells were transfected with GFP_2A_Cas9 and appropriate gRNA expression vectors at a DNA ratio of 1:1 (w:w). Two days after transfection cells were single cell sorted using a FACSJazz (BD Bioscience), gating for GFP positive cell population as described previously^46^. Indels in targeted genes were verified by Next Generation Sequencing (NGS) as described previously^46^. Primers are listed in Supplementary Table S2. Three clones with a confirmed indel and two control clones without indels were and expanded to 30 mL media before they were frozen down at 1 × 10^7^ cells per vial in spent CD-CHO medium with 5 % DMSO (Sigma-Aldrich).

### Characterizing CHO-S^Cas9^ functionality

To characterize Cas9 functionality we transfected clonal CHO-S^Cas9^ cells with a vector expressing gRNA against Mgat1 and verified indel generation on a pool level by NGS as described previously^48^ (using gRNA oligo primers MGAT1_gRNA_fwd and MGAT1_gRNA_rev and NGS primers MGAT1_miseq_fwd and MGAT1_miseq_rev listed in Supplementary Table S2). To analyze GFP expression, clonal cells were seeded in wells of a 96-well optical-bottom microplate (Greiner Bio-One) and identified GFP positive cells on the Celigo Cell Imaging Cytometer (Nexcelom Bioscience) using the green fluorescence channel. GFP negative gating was set on the basis of fluorescence emitted from CHO-S wild type cells.

### Library design and construction

For design of the metabolic gRNA library, a list of metabolic genes was extracted from the CHO metabolic network reconstruction^3^ along with a list of genes with metabolic GO terms in CHO and associated transcription factors. The gRNA templates were computationally designed using CRISPy (http://crispy.biosustain.dtu.dk/), resulting in a gRNA library with a minimum of 5 gRNAs per gene. The oligo library was synthesized by CustomArray. Full-length oligonucleotides were amplified by PCR using KAPA Hifi (Kapa Biosystems), size selected on a 2% agarose gel and purified with a QIAquick Gel Extraction Kit (Qiagen) as per manufacturer’s protocol. The gRNA-LGP vector (Addgene #52963) was digested using BsmBI (New England BioLabs) (4 µg gRNA-LGP vector, 5 µL buffer 3.1, 5 µL 10 x BSA, 3 µL BsmBI and H2O up to 50 µL were mixed and incubated at 55°C for 3 hours). Subsequently, 2 µL of calf intestinal alkaline phosphatase (New England BioLabs) was added to the digested vector and the mix was incubated at 37°C for 30 minutes before it was purified with a QIAquick PCR Purification Kit (Qiagen) as per manufacturer’s protocol. To assemble the gRNAs into the vector a 20 µL Gibson ligation reaction (New England BioLabs) was carried out (25 ng linearized vector, 10 ng purified insert, 10 µL 2 x Gibson Assembly Master Mix (New England BioLabs) and up to 20 µL H2O were mixed and incubated at 50°C for 1 hour). The assembled vector was purified using QIAquick PCR purification (Qiagen) and transformed into chemically competent E. coli (Invitrogen). Transformed bacteria were plated onto LB-carbenicillin plates for overnight incubation at 37°C, and plasmid DNA was purified using a HiSpeed Plasmid Maxi Kit (Qiagen).

### Lentiviral packaging

To produce the lentivirus, HEK293T cells were cultivated in DMEM supplemented with 10% Fetal Bovine Serum (FBS). One day prior to transfection, cells were seeded in a 15cm tissue culture plate at a density suitable for reaching 70-80% confluency at time of transfection. Culture medium was replaced with prewarmed DMEM containing 10% FBS. 36 µL Lipofectamine 3000 (Life Technologies) was diluted in 1.2 mL OptiMEM (LifeTechnologies) and in a separate tube 48 µL P3000 reagent, 12 µg pCMV (Addgene #12263), 3 µg pMD2.G (Addgene #12259) and 9 µg lentiviral vector were diluted in 1.2 mL OptiMEM. The solutions were incubated for 5 minutes at room temperature, mixed, incubated for another 30 minutes before they were added dropwise to the HEK293T cells. 48 hours and 72 hours after transfection the viral particles were concentrated using Centricon Plus-20 Centrifugal ultrafilters (100 kDa pore size), aliquoted and stored at −80°C.

### Puromycin kill curve

To determine the concentration of puromycin to be used to select the CHO library cells for gRNA insertion, a puromycin kill curve for CHO cells was determined. CHO-S wild type cells at a concentration of 1 × 10^6^ cells/mL in media containing various amounts of puromycin (0, 0.25, 0.5, 1, 2, 3, 4, 5, 6, 7, 8 and 10 µg/mL). Cell viability and VCD was monitored over 7 days and based on halted growth and complete cell death of wild type cells 10 µg/mL was used for further experiments (Supplementary Figure S3).

### Transducing CHO-S^Cas9^ with library virus

CHO-S^Cas9^ cells were seeded at 0.3 × 10^6^ cells/mL in 1 mL media in 26 wells of 12 well plates (BD Biosciences). In 25 of the wells, cells were transduced with 4 µL library virus/well along with 8 µg/mL Polybrene (Sigma-Aldrich) aiming for an MOI at 0.3-0.4 (Supplementary Methods and Results). Cells in the remaining well were left non-transduced as a negative control. After 24 hours, the cells were washed in PBS (Sigma-Aldrich) by centrifugation at 200 x g, resuspended in media and seeded in a new 12 well plate. After 24 hours, cells were expanded to 3 mL media in wells of 6 well plates (BD Biosciences). Selection for cells containing the gRNA insert was initiated by adding 10 µg/mL puromycin (Thermo Fisher Scientific) to each well (see puromycin kill curve in Supplementary Figure S3). Non-transduced control cells were monitored for complete cell death, equating finalised selection. The cells were washed and passed twice before they were expanded to attain enough cells to create a cell bank. Cells were frozen down at 1 × 10^7^ cells per vial in spent CD-CHO medium with 5 % DMSO (SigmaAldrich) and will from here on be referred to as CHO-S^Cas9^ library cells.

### Screening and DNA extraction

CHO-S^Cas9^ library cells were thawed in 30 mL media and expanded to 60 mL before starting the screen. On day 0 (T0) 1.5 × 10^7^ cells were spun down at 200 x g and resuspended in 60 mL appropriate screening media. The cells were grown for 14 days (passed to 0.25 × 10^6^ cells/mL every third day). 30 × 10^6^ cells were collected at T0 and on day 14 (T14). The pellets were stored at −80°C until further use. gDNA extraction of all 30 × 106 cells was carried out using GeneJET Genomic DNA Purification Kit (Thermo Fisher Scientific) following the manufacturer’s protocol. gDNA was eluted in 100 µL preheated elution buffer from the purification kit and incubated for 10 minutes before final centrifugation for maximum gDNA recovery.

### Preparation for next generation sequencing

50 µL PCR reactions with 3 µg input gDNA per reaction were run using Phusion® Hot Start II High-Fidelity DNA Polymerase (Thermo Fisher Scientific) (95°C for 4 min; 30 times: 98°C for 45s, 60°C for 30 s, 72°C for 1 min; 72°C for 7 min) using primers flanking the gRNA insert containing overhang sequenced compatible with Illumina Nextera XT indexing and 8 random nucleotides to increase the diversity of the sequences (LIB_8xN_NGS_FWD and LIB_8xN_NGS_REV listed in Supplemental Table S2). Double size selection was performed using Agencourt AMPure XP beads (Beckman Coulter) to exclude primer dimers and genomic DNA. The amplicons were indexed using Nextera XT Index Kit v2 (Illumina) sequence adapters using KAPA HiFi HotStart ReadyMix (KAPA Biosystems) (95°C for 3 min; 8 times: 95°C for 30s, 55°C for 30 s, 72°C for 30 s; 72°C for 5 min) and subjected to a second round of bead-based size exclusion. The resulting library was quantified with Qubit® using the dsDNA HS Assay Kit (Thermo Fisher Scientific) and the fragment size was determined using a 2100 Bioanalyzer Instrument (Agilent) before running the samples on a NextSeq 500 sequencer (Illumina).

### Analysis

Raw FASTQ files for samples from the end time points of glutamine selection were uploaded to PinAPL-PY (http://pinapl-py.ucsd.edu/)^49^ along with a file containing the sequences for all gRNAs contained in the library. Top candidates for enriched and for depleted gRNAs were ranked by an adjusted robust rank aggregation (aRRA) method^50^ and filtered for significance, compared between the replicates and used for verification of the screen. The screen was analyzed using default parameters set by PinAPL-PY.

### Batch culture

*Abhd11* knockout cell lines were seeded at 0.3 × 10^6^ cells/mL in 90 mL CD-CHO media with and without glutamine supplemented with 1 µl/mL AC in 250 mL Corning® Erlenmeyer culture flasks (Sigma-Aldrich). Cell viability and density were measured every day for a maximum of fourteen days.

### Analysis of cell line adapted to absence of glutamine by directed evolution

A previously established cell line that was adapted to grow without glutamine by stepwise decrease in glutamine concentration and directed evolution^32^ was grown in batch culture as previously described^51^. Samples were taken at the same time points and analysed using a mouse Agilent 22 k microarray (G4121B) platform as described for the parental cell line grown in medium with 8mM glutamine^51^. Differential transcriptome and statistical analyses were performed as previously described^51^.

## Supporting information

Supplementary Datafile 1

Supplementary Datafile 2

## Acknowledgements

The authors wish to thank Nachon Charanyanonda Petersen for assistance, cell line generation and batch culture and Anna Koza, Alexandra Hoffmeyer, Pannipa Pornpitapong for assistance with NGS, Dr. Prashant Mali for packaging the gRNA library into the lentivirus, Dr. James A Nathan for discussions regarding *Abhd11* and Daria Sergeeva for co-drawing Figure 1. This work was supported by the Novo Nordisk Foundation (NNF10CC1016517 and NNF16OC0021638) and NIGMS (R35 GM119850, NEL).

## ^1^ Abbreviations

αkgdhc: alpha ketoglutarate dehydrogenase complex
Cas9: CRISPR-associated protein 9
CHO: Chinese hamster ovary
CPM: counts per million
CRISPR: clustered regularly interspaced short palindromic repeats
DAPI: 4’,6-diamidino-2-phenylindole
GFP: green fluorescent protein
GLS: glutaminase
GLUL: glutamine synthetase
gRNA: guide RNA
Mgat1: mannosyl (alpha-1,3-)- glycoprotein beta-1,2-N-acetylglucosaminyltransferase
NGS: next generation sequencing
RNAi: RNA interference
TALEN: transcription activator-like effector nucleases
VCD: viable cell density
ZFN: zinc-finger nuclease

