## Supplementary Material for "A metabolic CRISPR-Cas9 screen in Chinese hamster ovary cells identifies glutamine-sensitive genes"

### Overview

**Supplementary methods and results**

Multiplicity of infection in CHO cells

Multiplicity of infection calculations

RNA extraction

Library preparation and RNASeq

RNASeq analysis

Transcriptomic analysis implicates attenuation of the interferon response with improved growth in glutamine free media

**Supplementary figures**

Supplementary Figure S1: Growth profile of the CHO-S$^{Cas9}$ CRISPR knockout screen

Supplementary Figure S2: Selection during screening

Supplementary Figure S3: Puromycin kill curve

Supplementary Figure S4: Differential expression of genes involved in the interferon response shows an attenuated response in *Abhd11* knockout cells when shifted to glutamine free medium

**Supplementary tables**

Supplementary Table S1: CHO-S$^{Cas9}$ validation by indel analysis

Supplementary Table S2: Primers and oligos

**Supplementary datafiles**

Supplementary Datafile 1: List of gRNAs in library

Supplementary Datafile 2: List of differentially expressed genes during adaptation to glutamine free media

### Supplementary methods and results

#### Multiplicity of infection in CHO cells

CHO-S wild type were seeded at 0.3 x $10^6$ cells/mL in 1 mL media in 5 wells of a 12 well plate (BD Biosciences). Four wells were transduced with 4, 10, 20 and 40μL virus/well along with 8 μg/mL Polybrene (Sigma-Aldrich). Cells in the remaining well were left non-transduced as a negative control. After 24 hours, the cells were washed in PBS (Sigma-Aldrich) by centrifugation at 200 x g, resuspended in media and seeded in a new 12 well plate. After 24 hours cells were expanded to 3 mL media in wells of 6 well plates (BD Biosciences). Selection for cells containing the gRNA insert was initiated by adding 10 μg/mL puromycin (Thermo Fisher Scientific) to each well. Non-transduced control cells were monitored for complete cell death, consonant with finalised selection. After selection, single transduced cells were deposited into Corning® 384 well plates (Sigma-Aldrich) containing 30 μL media without AC supplemented with 1% antibiotic/antimycotic (Gibco), and 1.5% HEPES buffer (Gibco). After 14 days of static incubation, viable cells were transferred to an MD96F Falcon™ plate (Thermo Fisher Scientific). Once confluent, 50 uL cell suspension from each well was transferred to a MicroAmp Fast 96 well reaction plate (Thermo Fisher Scientific). The plate was centrifuged at 1000 x g for 10 minutes, then the supernatant was removed via rapid inversion. Genomic DNA (gDNA) was extracted by resuspending the pellet in 20 μL QuickExtract™ DNA Extraction Solution (Epicentre) and processing the plate in the thermocycler (65°C for 15 minutes followed by 95°C for 5 minutes). gRNA integration was verified by NGS using a modified version of the Illumina 16S Metagenomic Sequencing

Library Preparation as described previously[1] (using primers lib_miseq_fwd, lib_miseq_rev, OMA1_fwd and OMA1_rev listed in Supplementary Table S3).

**Multiplicity of infection calculations**

When creating a CRISPR-Cas9 library using lentiviruses, the initial number of infections per cell, and thus number of gRNA per cell, follows a Poisson distribution[2]. That distribution depends on the ratio of viruses to cells. This is known as the "multiplicity of infection" or MOI. In CRISPR-Cas9 screens it is important to transduce at a low MOI to decrease the number gRNA integration events per cell. If a cell carries multiple different gRNAs the genotype to phenotype relation becomes complex to analyze. This confounder is alleviated by having multiple different gRNAs targeting each gene. However, a low MOI also increases the number of cells needed for 100% library coverage which can be a practical limitation. Most CRISPR-Cas9 screens report transduction with an MOI of 0.2-0.5[3–12]. Though a reason is not always given, one can speculate that the reason is that at an MOI of ~0.3, the Poisson distribution probability mass function (Poisson pmf) predicts that the fraction of cells having more than 1 gRNA should be:

$$p(k) = \frac{\lambda^k e^{-\lambda}}{k!} \Rightarrow p(k > 1) = 1 - p(k = 0) - p(k = 1) = 1 - e^{-\lambda} - \lambda e^{-\lambda}$$

where p(k) is the fraction of cells with k infections and lambda is the MOI. So with MOI = 0.3, the fraction of cells with more than 1 infection, p(k>1), will most likely be ~3.7%, which presumably is then considered sufficiently low to prevent analysis issues.

However these papers often seem to lack any description as to how they actually obtain this MOI and/or measure it[4,6,10,13–24]. Furthermore, if they obtain/measure this MOI before antibiotic selection against uninfected cells (wild type cells), then the fraction of cells with more than 1 insertion in the final library will be much higher than 3.7% because all the uninfected cells are ideally gone. With the uninfected cells (k=0) gone, the formula changes to:

$$p_{after}(k) = \frac{\left(\frac{\lambda^k e^{-\lambda}}{k!}\right)}{p(k>0)} \Rightarrow \frac{\left(\frac{\lambda^k e^{-\lambda}}{k!}\right)}{1-e^{-\lambda}} \Rightarrow \frac{\lambda^k}{(e^{\lambda}-1)k!}$$

where $p_{after}$ is the fraction of cells with k infections **after** antibiotic selection and k is larger than 0. So if you had an MOI of 0.3 before selection, then the fraction of cells with more than 1 infection after antibiotic selection, $p_{after}(k>1)$, would most likely be ~14%. We were only able to identify 1 other study that explicitly states this[25] and only 1 study that have selected an MOI that would lead to $p_{after}(k>1)$ being ~2.5%[22].

In this study we decided to aim for the lowest MOI possible where we could practically handle the number of cells needed for ~100% coverage of the library. Most of the previously referenced CRISPR screen studies estimate MOI by infecting a cell population with their lentiviral library and then counting the number of cells surviving with vs without an antibiotic selection. In such a study you are essentially counting the number of uninfected cells and assuming that the number of gRNAs per cell will otherwise follow the Poisson distribution. We speculate that it is possible that cells with multiple infections, and thus multiple antibiotic resistance genes, might survive better and thus perturb the assumed distribution. For this reason, we decided to count the number of different gRNAs per cell after antibiotic selection.

We set up an experiment identical to the process used in the actual library creation but with four different volumes of the viral library, 4, 10, 20 and 40 μL. After antibiotic selection, we single cell sorted the four resulting libraries, harvested gDNA from each clonal population and sequenced the gRNA insert and a control gene, OMA1. This allowed us to count how many different gRNA we had in each cell. We could then directly count how many percent of the cells had more than 1 gRNA and fitting this data to the $p_{after}(k)$ function, we could estimate lamda (pre-selection MOI) and estimate confidence intervals:

| μL virus | Percent having more than 1 gRNA (lower/upper 95% conf interval) | Fitted lamda (moi) (lower/upper 95% conf interval) |
|---|---|---|
| 4 | 11.8% [7.3%, 17.0%] | 0.25 [0.15, 0.36] |
| 10 | 30.1% [22.9%, 37.3%] | 0.68 [0.5, 0.87] |
| 20 | 35.3% [28.2%, 42.2%] | 0.81 [0.63, 1.01] |
| 40 | 56.8% [49.4%, 63.8%] | 1.50 [1.24, 1.78] |

Based on the above results, we chose to use 4 μL virus per 0.6 x $10^6$ cells/mL for transduction.

**RNA extraction**

During batch culture 2 x $10^6$ cells were harvested in mid-exponential phase. Cells were spun down at 200 x g, supernatant was discarded and the cell pellet was resuspended in 600 µL TRIzol™ Reagent (Thermo Fisher Scientific). RNA was isolated using Direct-zol™ RNA MiniPrep Plus (Zymo Research) following manufacturer's protocol. RNA concentration was measured with Qubit fluorometric analysis (Life Technologies) and RNA quality was determined with Agilent 2100 bioanalyzer and Fragment analyzer automated CE system (Advanced Analytical Technologies, Inc.).

**Library preparation and RNASeq**

The RNA samples were processed by the NGS lab at the Novo Nordisk Foundation Center for Biosustainability (Technical University of Denmark). The samples were prepared with Illumina's TruSeq Stranded mRNA sample preparation kit according to manufacturer's instructions, pooled and sequenced on a NextSeq 500 machine (Illumina) using the NextSeq High Output Kit v2, for 75 cycles of single-end reads for an average of 10 million reads per sample. Raw reads deposited in the SRA (BioProject: PRJNA630824).

**RNASeq analysis**

Sequencing reads were aligned against the NCBI *C. griseus* "PICR" genome[26] (GCF_003668045.1) using STAR[27]. Aligned reads were quantified using HTSeq[28] to obtain counts for annotated genes. Differential gene expression was determined by DESeq2[29] following their standard workflow. Comparisons were made for *Abhd11* knockout vs control cells in media containing and lacking glutamine as well as *Abhd11* knockout or control cells in media lacking glutamine vs control cells in media containing glutamine. CHO gene IDs were mapped to human genes and analyzed using GSEA[30]. Three specific gene sets, Hallmark

Interferon Alpha Response, Hallmark Interferon Gamma Response, and Browne Interferon Responsive Genes[31] were examined on the individual gene level.

**Transcriptomic analysis implicates attenuation of the interferon response with improved growth in glutamine free media**

Gene set enrichment analysis of a comparison between the control cells in mid-exponential phase in media with and without glutamine showed significant enrichment in numerous interferon associated gene sets (data not shown), which was surprising given their canonical association with viral infection. However, as some of the interferon stimulated responses leads to increased RNAse L activity and translational inhibitaion via eIF2α phosphorylation[32], this could be a reason for retarded growth in media lacking glutamine. Interestingly, not only does growth in glutamine free medium induce a weaker activation of interferon responsive genes in knockout cells than wildtype cells (compared to the wildtype cell grown in media with glutamine), the knockout cells also have fewer of these genes differential expressed (Supplementary Figure S4). The directionality of this relationship is uncertain, specifically it is unclear if glutamine deprivation induces the interferon response leading to a decrease in growth or if glutamine deprivation leads to a decrease in growth that induces the interferon response due to a misinterpretation of biochemical cues. However, while the mechanisms are unclear, the effect is striking.

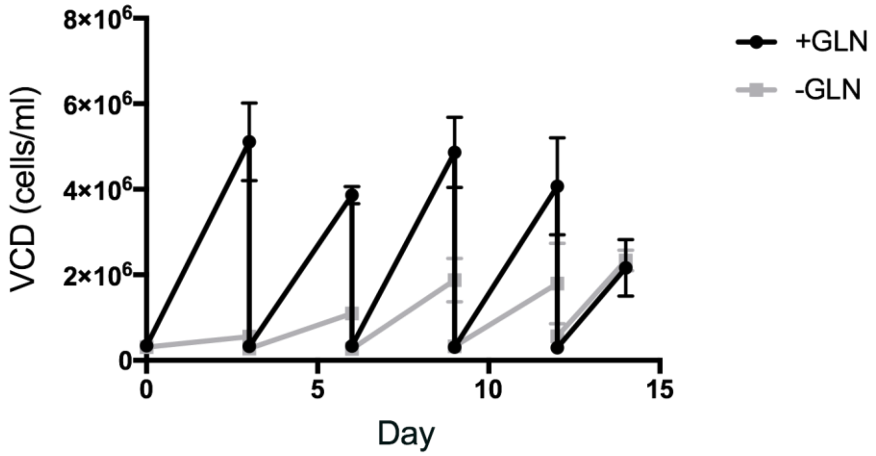


**Supplementary Figure S1: Growth profile of the CHO-S$^{Cas9}$ CRISPR Knockout Screen**

CHO-S$^{Cas9}$ CRISPR KO screens were grown for 14 days in media supplemented with (black) and without (grey) glutamine (GLN) (n=3 for both conditions). Cells were passed and viable cell density (VCD) was measured every 3 days.

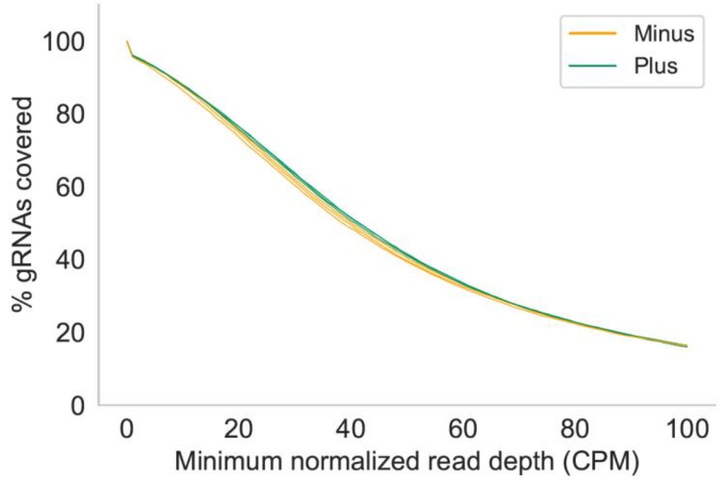


**Supplementary Figure S2: Glutamine depletion acts as a weak selective pressure**

Percentage of gRNAs with a minimum normalized read depth is shown for cells grown in medium with (green) and without (orange) glutamine for 14 days (n=3, both conditions). There is little difference between the gRNA distribution, in agreement with the weakness of glutamine as a selective pressure.

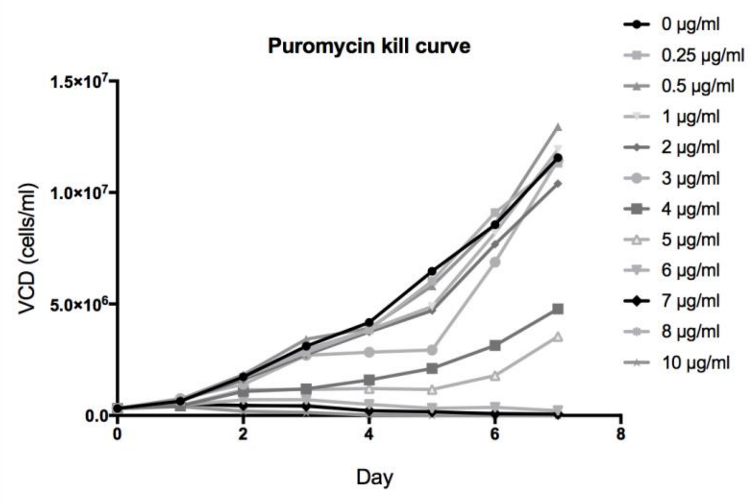


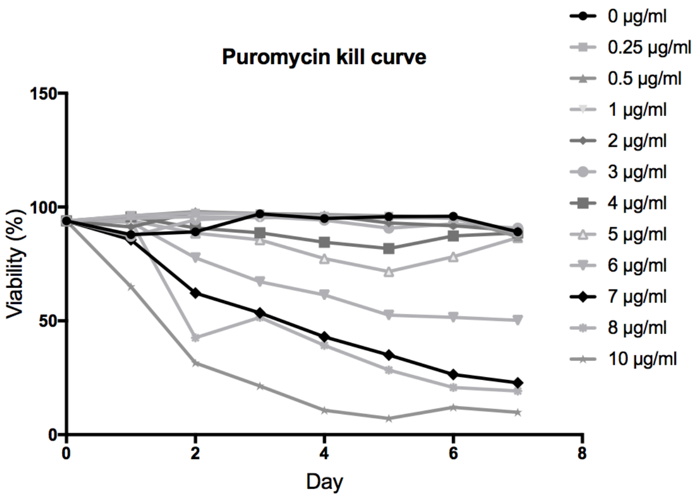


**Supplementary Figure S3: Puromycin Kill Curve**

CHO-S cells were treated with varying concentrations of puromycin (spanning 0 – 10 μg/mL). Viable cell density (VCD) and viability were measured for 7 days. We continued to work with a concentration of 10 μg/mL for sufficient stagnant VCD and decrease in viability.

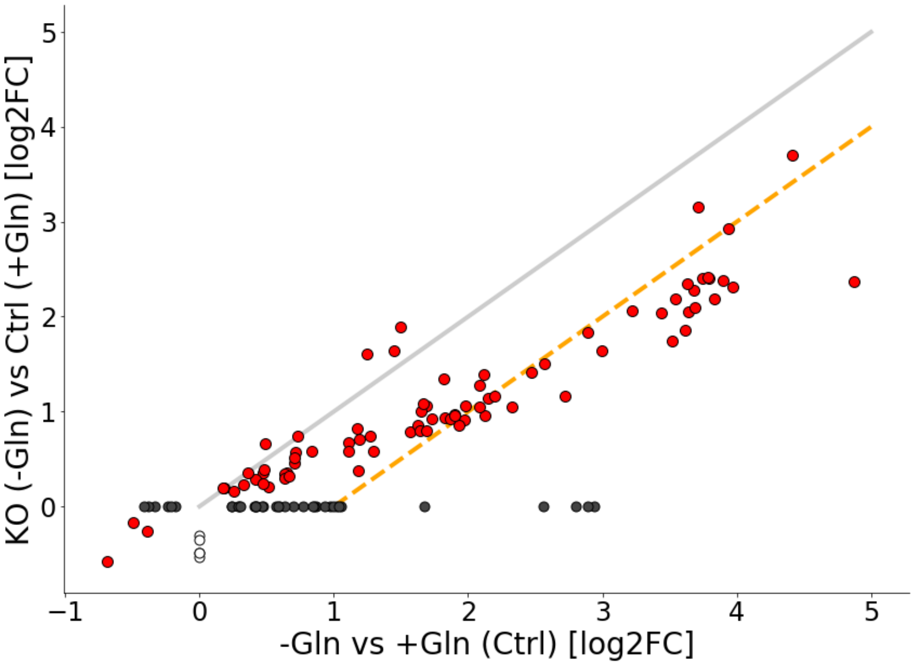


**Supplementary Figure S4: Differential expression of genes involved in the interferon response shows an attenuated response in *Abhd11* knockout cells when shifted to glutamine free medium**

Log 2 fold changes in differentially expressed interferon response genes are compared between two comparisons: The x-axis shows log2 fold changes for the comparison between the control cell grown with and without glutamine while the y-axis shows the comparison between the knockout cell line grown without glutamine and the control cell line grown with glutamine. Genes shown in red (n=80) are differentially expressed in both comparisons. Genes in white (n=5) are differentially expressed only in the knockout vs control comparison and genes in dark gray (n=37) are differentially expressed only in the control vs control comparison. The gray line shows y=x, the orange line y=x-1 (i.e., log2 fold change is 1 less in the knockout vs control comparison than in the control vs control comparison).

### Supplementary tables

#### Supplementary Table S1: CHO-S$^{Cas9}$ Validation by Indel Analysis

CHO-S$^{Cas9}$ cells were transfected with gRNA targeting the Mgat1 gene and sampled for indel analysis at a pool level. The above indel scores are of 5 different clonal CHO-S$^{Cas9}$ cell lines from two different experiments. Cas9 showed ~50-70% cutting efficiency for all clonal cell lines.

| Sample | Target | %Indel |
|---|---|---|
| CHO-S$^{Cas9}$_L1_mGat1 | Mgat1 | 66,81% |
| CHO-S$^{Cas9}$_L15_mGat1 | Mgat1 | 69,16% |
| CHO-S$^{Cas9}$_L21_mGat1 | Mgat1 | 59,12% |
| CHO-S$^{Cas9}$_positive control | Mgat1 | 69,19% |
| CHO-S$^{Cas9}$_negative control | Mgat1 | 0,44% |

| Sample | Target | %Indel |
|---|---|---|
| CHO-S$^{Cas9}$_L12_mGat1 | Mgat1 | 51,93% |
| CHO-S$^{Cas9}$_positive control | Mgat1 | 67,63% |
| CHO-S$^{Cas9}$_negative control | Mgat1 | 0,23% |

### Supplementary Table S4: Primers and Oligos

| Primer/oligo name | Sequence (Green: NGS compatible overhangs) |
|---|---|
| MGAT1_gRNA_fwd | GGAAAGGACGAAACACCGTGGAGTTGGAGCGGCAGCGGTTTTAGAGCTAGAAAT |
| MGAT1_gRNA_rev | CTAAAACCGCTGCCGCTCCAACTCCACGGTGTTTCGTCCTTTCCACAAGATAT |
| MGAT1_miseq_fwd | TCGTCGGCAGCGTCAGATGTGTATAAGAGACAGTTCTGGACACGCCCAGC |
| MGAT1_miseq_fwd | GTCTCGTGGGCTCGGAGATGTGTATAAGAGACAGGCCACGGTGGGCACTTT |
| Lib_miseq_fwd | TCGTCGGCAGCGTCAGATGTGTATAAGAGACAGATCTTGTGGAAAGGACGAAACAC |
| Lib_miseq_rev | GTCTCGTGGGCTCGGAGATGTGTATAAGAGACAGCTCGGTGCCACTTTTTCAAGTT |
| OMA1_fwd | TCGTCGGCAGCGTCAGATGTGTATAAGAGACAGTCCCATGAAGCAGCATAAGCA |
| OMA1_rev | GTCTCGTGGGCTCGGAGATGTGTATAAGAGACAGAGAGATCCCTGGGACATCCT |
| LIB_8xN_NGS_FWD | TCGTCGGCAGCGTCAGATGTGTATAAGAGACAGNNNNNNNNGGCTTTATATATCTTGTGGAAAGGACGAAACACC |
| LIB_8xN_NGS_REV | GTCTCGTGGGCTCGGAGATGTGTATAAGAGACAGNNNNNNNNCCGACTCGGTGCCACTTTTTCAA |
| ABHD11_gRNAfwd | GGAAAGGACGAAACACCGAGCGTGGAGAGTCCCCCGTGTTTTAGAGCTAGAAAT |
| ABHD11_gRNArev | CTAAAACACGGGGGACTCTCCACGCTCGGTGTTTCGTCCTTTCCACAAGATAT |
| ABHD11_Miseqfwd | TCGTCGGCAGCGTCAGATGTGTATAAGAGACAGCAGAGCAGGAAGATAGGCGG |
| ABHD11_Miseqrev | GTCTCGTGGGCTCGGAGATGTGTATAAGAGACAGGGTGTCCTTGACTGACCTCG |
